# Solid-state NMR spectroscopy identifies three classes of lipids in *C. neoformans* melanized cell walls and whole fungal cells

**DOI:** 10.1101/2020.07.13.200741

**Authors:** Christine Chrissian, Emma Camacho, John E. Kelly, Hsin Wang, Arturo Casadevall, Ruth E. Stark

## Abstract

A primary virulence-associated trait of the opportunistic fungal pathogen *Cryptococcus neoformans* is the production of melanin pigments that are deposited into the cell wall and interfere with the host immune response. Previously, our solid-state NMR studies of isolated melanized cell walls (melanin ‘ghosts’) revealed that the pigments are strongly associated with lipids, but their identities, origins, and potential roles were undetermined. Herein, we exploited spectral editing techniques to identify and quantify the lipid molecules associated with pigments in melanin ghosts. The lipid profiles were remarkably similar in whole *C. neoformans* cells, grown under either melanizing or non-melanizing conditions; triglycerides (TGs), sterol esters (SEs) and polyisoprenoids (PPs) were the major constituents. Although no quantitative differences were found between melanized and non-melanized cells, melanin ghosts were relatively enriched in SEs and PPs. In contrast to lipid structures reported during early stages of fungal growth in nutrient-rich media, variants found herein could be linked to nutrient stress, cell aging, and subsequent production of substances that promote chronic fungal infections. The fact that TGs and SEs are the typical cargo of lipid droplets suggests that these organelles could be connected to *C. neoformans* melanin synthesis. Moreover, the discovery of PPs is intriguing because dolichol is a well-established constituent of human neuromelanin. The presence of these lipid species even in non-melanized cells suggests that they could be produced constitutively under stress conditions in anticipation of melanin synthesis. These findings demonstrate that *C. neoformans* lipids are more varied compositionally and functionally than previously recognized.

## Introduction

*Cryptococcus neoformans* is a globally distributed opportunistic fungal pathogen and the primary causative agent of cryptococcosis, a frequently lethal infectious disease that results in approximately 180,000 deaths each year worldwide (1). Avoiding contact with *C. neoformans* is essentially impossible due to its global presence and ability to thrive in soil, trees, and bird droppings, among other environmental niches. Initial exposure is thought to occur during early childhood through the inhalation of airborne spores or desiccated yeast cells, usually leading to asymptomatic pulmonary infection but no clinical manifestation of disease (2). In immunocompetent hosts, the infection is either eliminated or becomes latent. However, in immunocompromised individuals, the outcome is typically cryptococcal pneumonia followed by rapid dissemination to the central nervous system (3). Ultimately the result is meningitis, which is both the most commonly presented and most life-threatening form of cryptococcosis. Cryptococcal meningitis is responsible for approximately 15% of all HIV/AIDS-related deaths annually (1). The efficacy of antifungal therapies against *C. neoformans* is also limited: even after intervention the mortality rate is approximately 50%, rendering the chance of survival comparable to that of a coin toss (4).

A primary virulence-associated trait in *C. neoformans* is the production of melanin pigments, which are deposited within the cell wall and form a defensive barrier to external stressors. Melanins are a group of biologically ubiquitous and structurally heterogeneous pigments produced via the oxidation and polymerization of particular aromatic ring compounds. They share a multitude of remarkable properties that include: the ability to absorb ultraviolet (UV) light, transduce electricity, and protect against ionizing radiation. In fungi, melanin pigments contribute to virulence by reducing cellular susceptibility to oxidative damage, inhibiting phagocytosis, and interfering with the host immune response to infection (5). *C. neoformans* is unique among melanotic fungi because of its dependence on exogenous substrates for melanin production; this organism is unable to synthesize melanin precursors endogenously and therefore must scavenge catecholamines and/or other phenolic compounds from the environment (6). *C. neoformans* cells isolated from individuals with active cryptococcal infections display robust pigment production (7–9), an unsurprising outcome considering that melanization is correlated with dissemination of infections and resistance to anti-fungal therapeutics (10, 11). For these reasons, the study of *C. neoformans* melanogenesis can contribute directly to our understanding of the pathogenicity of this organism.

The current model for *C. neoformans* melanization is a process in which intracellularly synthesized melanin pigments are extruded into the cell wall, where they then form strong associations with various molecular constituents that are essential for their deposition and retention (12). The best-established components of this melanization scaffold are the polysaccharides chitin and chitosan (13); for instance, *C. neoformans* cells that are unable to produce cell-wall chitin and chitosan display a ‘leaky melanin’ phenotype, in which melanin pigments are unable to anchor to the cell wall and consequently leak out into the culture media (14, 15). Previously, we used solid-state nuclear magnetic resonance spectroscopy (ssNMR) to study the molecular structure of melanized fungal cell walls in ‘melanin ghost’ preparations from which other cellular constituents were removed (16), confirming that melanin pigments are indeed attached to these polysaccharides (17). Even though the protocol used to prepare melanin ghosts is both enzymatically and chemically degrading, a portion of the cell wall that participates in melanin deposition survives this process. Moreover, chitin and chitosan are not the only cellular constituents that associate tightly with melanin pigments — a substantial proportion of the non-pigment moieties found in melanin ghosts have been shown to be lipids, with largely unknown identities, origins, and physiological roles (13).

In the current study, we aimed to 1) identify the lipid molecular species that associate with melanin pigments; and 2) determine whether melanized cells have a distinct lipid profile compared with non-melanized cells. As an alternative to attempting extraction of the strongly bound pigment-associated lipids from the melanin ghosts, we devised a ssNMR approach to characterize the chemical structures of these components in near-native cellular systems. Because *C. neoformans* cannot utilize endogenous substrates for melanin synthesis, cells grown in media containing [U-^13^C_6_]-glucose as the sole carbon nutrient source and natural abundance L-3,4-dihydroxyphenylalanine (L-DOPA) as the obligatory melanization precursor yielded melanized samples in which only the non-pigment cellular constituents are enriched in the NMR-active ^13^C isotope. Moreover, the requirement of *C. neoformans* for exogenous substrates to synthesize melanin pigments enabled us to compare the lipid profile of whole intact cells under melanizing and non-melanizing conditions. By leveraging the capabilities of NMR to select constituents that undergo a certain degree of molecular motion in the solid state, it was also possible to edit out ^13^C resonances from rigid polysaccharides so that unambiguous peak assignments of the mobile lipid species could be made.

## Results

### 2D ^13^C-^13^C and ^1^H-^13^C experiments reveal sterol esters, triglycerides, and polyisoprenoids as the major lipid constituents in melanin ghosts

Lipid constituents were first identified in *C. neoformans* melanin ghosts by our group in 2008 (18). In that work, high-resolution magic-angle spinning (HRMAS) NMR experiments conducted on melanin ghosts swelled with DMSO revealed ^1^H-^13^C through-bond connectivities that were surprisingly consistent with triglycerides (TGs). Considering that melanin ghosts are isolated using exhaustive chemical degradation, including three successive lipid extraction steps and boiling in strong acid, the finding of TGs seemed counterintuitive. Nevertheless, our follow-up studies have corroborated this result, and moreover, they have suggested that other lipid species in addition to TGs also resist extraction and are retained in the ghosts (13).

A noteworthy feature of these lipids is that, despite their strong association with melanin pigments, they exhibit a significant degree of molecular mobility in the solid state and effectively undergo rapid isotropic reorientation even in dry preparations. In addition to raising an intriguing question regarding the molecular interactions that govern melanin pigment-lipid attachment, this retention of molecular mobility materially improved our ability to identify the individual lipid components of melanin ghosts using ssNMR. Thus, it first allowed us to implement spectral editing techniques that discriminate between constituents based on variations in molecular motion: by preferentially detecting the signals arising from the mobile lipids and filtering out resonances from relatively rigid moieties such as polysaccharides, we were able to alleviate much of the spectral overlap that can compromise structural analyses in macromolecular biological systems. Second, molecules that undergo isotropic reorientation gave rise to sharp NMR signals, because motional averaging abolishes through-space ^1^H-^13^C dipolar couplings that would otherwise be a primary source of spectral line broadening. As a result, we successfully acquired high-resolution ^13^C NMR spectra that displayed few overlapping signals and full width at half maximum linewidths less than 1 ppm, facilitating the unambiguous assignment of all observed lipid resonances.

To characterize the molecular structures of the pigment-associated lipids, we conducted a set of 2D experiments that measured ^13^C-^13^C and ^1^H-^13^C through-bond connectivities, respectively. Correlations between directly bonded pairs of ^13^C-^13^C nuclei were obtained by implementing a 2D J-INADEQUATE experiment that was carried out with direct polarization (DP) and a short (2-s) recycle delay for ^13^C signal acquisition in order to favor the signals from mobile constituents (19). In 2D J-INADEQUATE spectra, a pair of covalently bonded carbons is represented by two cross-peaks that each have unique single quantum (SQ) ^13^C chemical shifts but share a common double quantum (DQ) ^13^C chemical shift. The DQ shift is the sum of the two SQ shifts for a directly bonded carbon pair, so signal overlap is reduced by spreading the SQ shifts over a larger frequency range. To complement these data and verify our peak assignments, we also carried out a 2D ^1^H-^13^C HETCOR experiment using INEPT as the polarization transfer step, obtaining crosspeaks for each bonded ^1^H-^13^C pair within an isotropically tumbling molecular fragment along with excellent sensitivity and spectral resolution (20).

Cursory inspection of the 2D DP J-INADEQUATE spectrum of melanin ghosts (Figure 1, top) revealed a significant number of resonances distributed over a wide chemical shift range, which suggested that melanin pigments are indeed associated with multiple lipid species, including TGs. Site-specific peak assignments for these molecules are summarized in Table S1. The presence of TGs, for which ^13^C NMR assignments are well documented (21–23), was immediately apparent from the well-resolved pair of SQ signals at 62 and 69 ppm that are correlated with the DQ sum of their SQ frequencies at 131 ppm. These correlations are consistent with a glycerol backbone in which each of the three carbons is esterified to a similar type of FA, resulting in a symmetrical molecule that gives rise to just two inequivalent resonances for the CHO and two CH_2_O groups, respectively. Cross-peaks characteristic of long-chain FAs were also readily observed: between C1-C2 (172 and 35 ppm), C16-C17 (32 and 23 ppm), and C17-C18 (23 and 14 ppm) (24, 25). The signals at 130 and 27 ppm correspond to a linkage between the olefinic and allylic carbons of unsaturated FAs; the allylic carbons at 27 ppm are linked in turn to chain methylene carbons resonating at 30 ppm (26, 27). Moreover, the correlation between signals at 129 and 26 ppm indicates a linkage between olefinic and bis-allylic carbons, suggesting that a substantial portion of the FAs present in melanin ghosts possess more than one point of unsaturation (26, 27).

**Figure 1.**
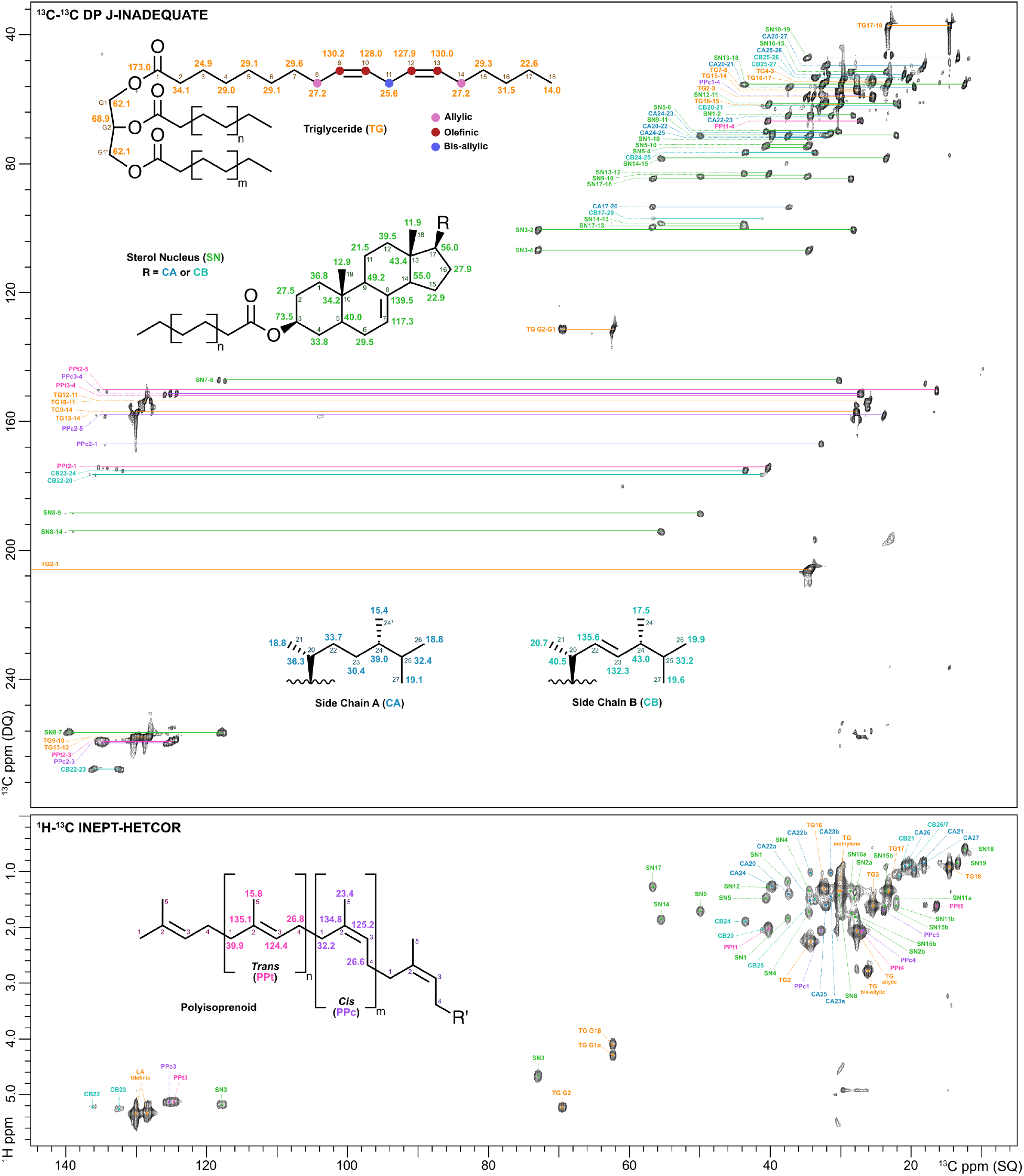
2D ^13^C-^1^3C DP J-INADEQUATE (top) and 2D ^1^H-^13^C INEPT-HETCOR (bottom) contour plots for *C. neoformans* melanin ghosts generated from H99 cells grown in [U-^13^C_6_]-glucose-enriched media. The displayed signals are attributed to triglycerides with unsaturated fatty acid acyl chains (**TG**; orange), the Δ7-sterol esters ergosta-7-enol and ergosta-7,22-dienol that share the same nucleus (**SN**; green) but have different side chains, designated as Chain A (**CA**; blue) and Chain B (**CB**; turquoise), respectively, and polyisoprenoids containing isoprene units in both the *cis* (**PPc**; purple) and *trans* (**PPt**; pink) configurations.

The 2D HETCOR experiment revealed additional ^1^H-^13^C bonded pairs that confirmed our TG structural identification (Figure 1, bottom). For instance, the carbon at 62 ppm that we attributed to the CH_2_O group of glycerol was found to correlate with two protons (4.07, 4.28 ppm), whereas the CHO carbon at 69 ppm was only correlated with one proton (5.21 ppm) (28–30). Each of these protons has chemical shifts consistent with TGs; ^1^H-^13^C cross-peaks corresponding to other possible types of esterified glycerol moieties (e.g., diglycerides) were not observed. The melanin ghost HETCOR spectrum also provided verification for the presence of di-unsaturated FAs: the bis-allylic carbon we identified in the J-INADEQUATE spectrum (26 ppm) is flanked by two double bonds, and consequently, the proton to which it is covalently bonded gives rise to a relatively downfield characteristic signal at 2.77 ppm (28–30). Nevertheless, completion of the TG resonance assignments left the majority of the resonances in both the ^13^C-^13^C J-INADEQUATE and ^1^H-^13^C HETCOR spectra unaccounted for.

Further analysis of these two 2D spectra revealed sterol esters (SEs) as a second major lipid component present within melanin ghosts. The ^13^C-^13^C through-bond information provided by the J-INADEQUATE experiment, in conjunction with the exquisite sensitivity of NMR chemical shifts to variations in molecular structure, enabled us to unambiguously identify two predominant sterols. Each sterol ester species had a structurally identical steroid nucleus with a single unsaturated linkage between C7 and C8 (118 and 139 ppm; Δ^7^ sterol) (31), but they differed in their sidechain structures. The C17 carbon that forms a covalent bond between the sterol nucleus and C20 of the sidechain displayed four cross-peaks, of which only two could be accounted for by linkages to adjacent carbons of the nucleus (C17-C13: 56 and 43 ppm; C17-C16: 56 and 28 ppm). Thus, the other cross-peaks involving the C17 carbon can be attributed to linkages with the C20 carbons of two chemically distinct sidechains. Although both sidechains were determined to share the same 24-methyl carbon backbone, one sidechain is fully saturated whereas the other contains a double bond between C22 and C23 (136 and 132 ppm). Distinguishing between these two sidechains allowed us to identify the predominant sterols in melanin ghosts as ergosta-7-enol and ergosta-7,22-dienol (31–36). Interestingly, our data strongly suggest that both sterol species are present only in their esterified forms. In free sterols, the chemical shifts of the hydroxylated C3 carbon and the hydrogen atom to which it is covalently bound would be 71 and 3.60 ppm, respectively. However, the HETCOR spectrum of melanin ghosts displayed a single peak in that chemical shift region with ^13^C and ^1^H values of 73 and 4.67 ppm, consistent with a compound in which the hydroxyl group is esterified (31, 33), likely to a long-chain FA.

The third and final group of lipid constituents that we identified in melanin ghosts are polyisoprenoids (PPs), which are linear polymers of isoprene units. Of the few remaining ^13^C-^13^C cross-peaks in the J-INADEQUATE spectrum that had not been attributed to TGs or SEs, of particular note were the correlations between a carbon at ~135 ppm and five different carbons at 16, 24, 32, 40 and 125 ppm. This observation suggested that the molecule contains two structurally similar carbons with overlapping resonances at 135 ppm but linkages to magnetically distinct repeating units. PPs fit this description, and their reported ^13^C and ^1^H chemical shifts (37–40) are consistent with our data. The repeating isoprene units that comprise PPs can adopt either *cis* or *trans* configurations, yielding distinct chemical shifts for the C1 and C5 carbon signals of each stereoisomer. Thus, the C2-C3 and C3-C4 linkages of both isomers can be accounted for by the single pair of cross-peaks at (135 and 125) and (125 and 27) ppm, respectively. Although the C2 carbon chemical shift does not differ between the two configurations, the C2, C1 and C2, C5 cross-peaks it forms are unique: those corresponding to the *cis* isomer were observed at (135 and 32) and (135 and 24) ppm, and to the *trans* isomer at (135 and 40) and (135 and 16) ppm. The presence of PPs in melanin ghosts is confirmed by our HETCOR data, which include atypically downfield ^1^H chemical shifts compared with most lipids and support substantial double bond content. Consequently, the ^1^H-^13^C pairs attributable to PPs are clearly visible because their cross-peaks are located in a relatively uncluttered spectral region. The protons covalently bonded to the C1 and C4 carbons of both *cis* and *trans* isomers were observed at ~2.0 ppm, and those bonded to the C5 carbons appeared at ~1.6 ppm (37, 38). Additionally, the ^1^H-^13^C cross-peak at (5.10, 125) ppm corresponded to the C3 carbon and its bonded proton, whereas the C2 carbon was not observed in a HETCOR experiment because it has no attached protons.

### Melanin ghosts, melanized whole cells, and non-melanized whole cells contain remarkably similar lipid species

The fact that the pigment-associated lipids we identified in melanin ghosts have not been reported previously as major constituents of *C. neoformans* cells led us to hypothesize that they might be produced exclusively under melanizing conditions. Our current understanding of *C. neoformans* lipid composition is based on only a handful of studies, all conducted on lipid extracts from cells grown in rich media under non-melanizing conditions (i.e., in the absence of an obligatory exogenous melanization precursor). The lack of lipid analyses for melanized cells does not necessarily reflect a lack of interest but could instead be attributed to experimental challenges: melanin pigments are exceptionally recalcitrant and thus melanized cells resist homogenization, which can diminish the effectiveness of lipid extractions. To circumvent this problem and determine whether melanized cells have a lipid profile distinct from non-melanized cells, we replicated the ssNMR strategy used to define the lipid profile of melanin ghosts to examine whole intact *C. neoformans* cells grown either with or without the melanization precursor L-DOPA.

In addition to using spectral editing techniques to select the NMR signals of mobile molecular moieties, the detection of the lipid constituents of interest was further enhanced by conducting our DP-INADEQUATE and INEPT-HETCOR experiments on acapsular cells from the Cap59 mutant strain. As an encapsulated yeast, approximately 50% of the *C. neoformans* cell mass is typically accounted for by a robust external layer of high molecular weight polysaccharides (41). Thus, the NMR response from this large proportion of capsular polysaccharides in wild-type (WT) H99 cells will present a dynamic range problem, overpowering the signals of constituents that are present in lesser quantities and compromising the detection of their spectral features (Figure S1). Conversely, the absence of capsular polysaccharides in Cap59 cells yields spectra that clearly display resonances attributable to a variety of cellular components (Figure 2).

**Figure 2.**
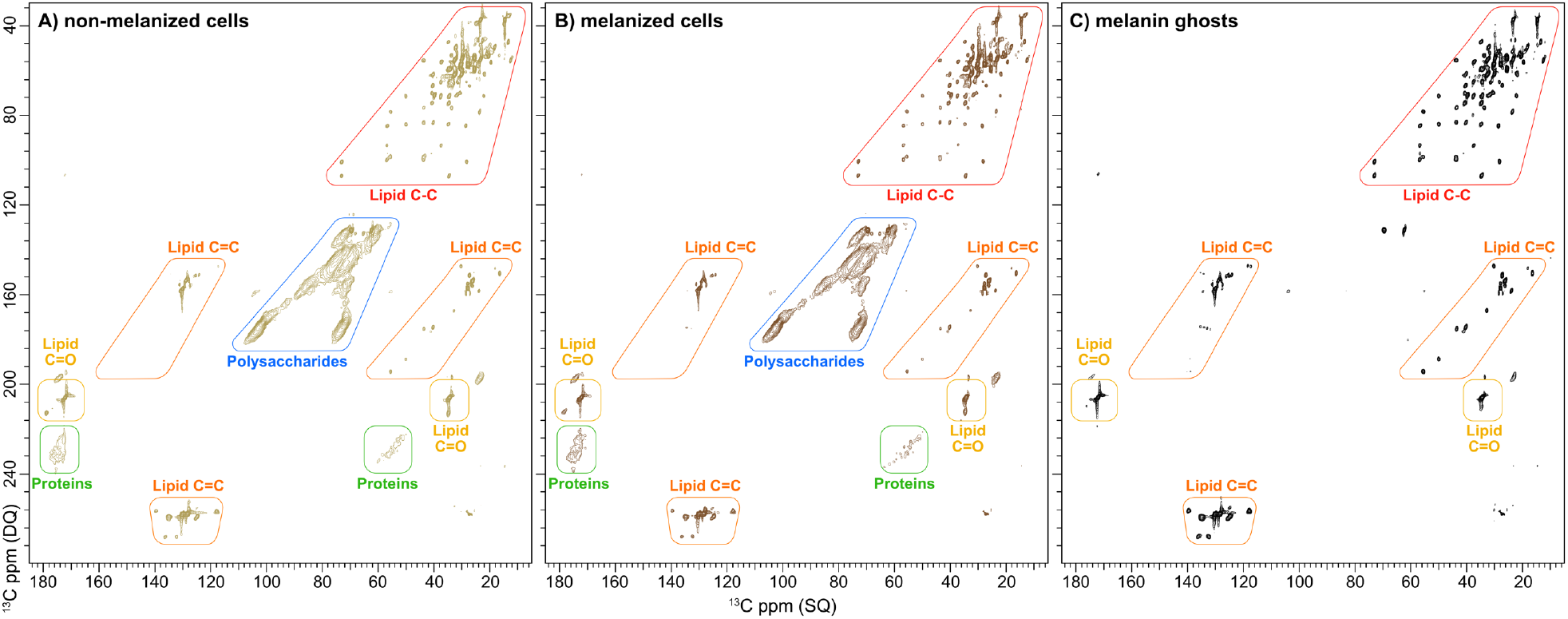
2D ^13^C-^13^C DP J-INADEQUATE spectra of *C. neoformans* whole cell and melanin ghost samples prepared from cultures containing [U-^13^C_6_]-glucose-enriched media. A) Non-melanized and B) melanized whole cells from the acapsular mutant Cap59 strain and C) melanin ghosts generated from H99 cells. The cross-peaks attributed to key functional groups have been demarcated to highlight the similarities between the three samples.

The 2D ^13^C-^13^C DP J-INADEQUATE spectra of non-melanized and melanized *C. neoformans* Cap59 whole cells (Figures 2A and 2B) and H99 melanin ghosts (Figure 2C) permitted us to compare their respective lipid compositions. Whereas only lipid signals are observed in the melanin ghost spectrum, the whole-cell spectra exhibited additional resonances; the majority of these features can be assigned to correlations between directly bonded ^13^C-^13^C pairs within the ring structures of non-chitinous cell-wall polysaccharides such as α- and β-1,3-glucans. As expected, these signals were absent from the DP J-INADEQUATE spectrum of melanin ghosts, because glucans are removed by enzymatic and chemical treatments during sample preparation. However, the finding that the DP J-INADEQUATE spectra of melanized and non-melanized cells are nearly indistinguishable was unanticipated. All of the well-resolved, and therefore identifiable, cross-peaks are clearly visible in both spectra (Figures 2A and 2B), demonstrating that melanization has no influence over the types of lipid species that are produced by *C. neoformans* cells. Even more surprising was our observation that both the DP J-INADEQUATE (Figures 2A and 2B) and INEPT-HETCOR (Figures S2A and S2B) spectra of each whole cell sample displayed identical lipid correlations to those displayed in the analogous spectra of melanin ghosts (Figures 1 and 2C). This finding indicates that the Δ^7^ SEs, TGs and PPs identified as major lipid components in melanin ghosts also comprise the lipid content of whole *C. neoformans* cells. Taken together, our data suggest that melanization does not trigger the synthesis of any particular lipid species, which is consistent with prior work that demonstrated melanization changes the transcriptional profiling of only a few genes in *C. neoformans* (42). Instead, it appears that *C. neoformans* melanin pigments associate with the lipids produced under non-melanizing conditions and due to the recalcitrant nature of the pigment, a significant portion of these lipids are able to resist the chemical and extractive treatments used for ghost preparation.

### Melanin ghosts have greater proportions of particular lipid species compared with whole melanized and non-melanized cells

Thus far we had identified the pigment-associated lipid species in melanin ghosts as Δ^7^ SEs, TGs and PPs, which we subsequently demonstrated are also the major species of lipids produced by *C. neoformans* cells under both melanizing and non-melanizing conditions. However, it was of further interest to determine whether the relative amounts of lipid constituents differ in melanized and non-melanized cells, and whether the proportions of Δ^7^ sterols, TGs and PPs are maintained between melanin ghosts and melanized whole cells. That is, we hypothesized that even if the same lipid species were present, the ease of their production or removal might be impacted by the presence of the melanin pigment.

To address these questions, we conducted direct polarization (DPMAS) ^13^C NMR experiments with a long recycle delay (50 s) to acquire spectra in which the integrated peak areas are quantitatively reliable and therefore reflect the relative amounts of the various cellular constituents in each sample. Despite the significant number of overlapping signals in these 1D spectra, we were able to locate several resonances that are unique to each lipid type using the carbon chemical shift information extracted from the 2D ^13^C-^13^C DP J-INADEQUATE and ^1^H-^13^C INEPT-HETCOR experiments described above. These identifications allowed us to carry out spectral deconvolution to determine the fraction of the total signal intensity attributable to each lipid constituent and thereby estimate its relative content in each of the *C. neoformans* samples.

Visual comparison of the ^13^C DPMAS spectra of non-melanized (Figure 3A) and melanized (Figure 3B) acapsular *C. neoformans* cells revealed no discernible quantitative differences in the relative amounts of lipids or other constituent types between the two samples: as noted in connection with our 2D J-INADEQUATE and HETCOR data, the 1D ^13^C DPMAS spectra of cells grown under melanizing and non-melanizing conditions were virtually indistinguishable from one another. Conversely, a number of striking visual differences were observed between the spectra of melanin ghosts (Figure 3C) and whole cells (Figures 3A and 3B); those trends were confirmed by the significant compositional variations revealed by our quantitative analyses. The clear dissimilarity between the two sample types is evidenced by the disparate relative contributions of lipids and polysaccharides to the overall signal intensity of each spectrum. Whereas signals attributable to polysaccharides (~58-108 ppm) dominate the whole cell spectra, accounting for approximately 68% of the overall signal intensity, the sum of the integrated peak areas corresponding to TGs, SEs, and PPs accounted for only 8% (Figure 3D). In contrast, for melanin ghosts the polysaccharide content was 2.5 times lower (27%), whereas the total lipid proportion was 5.3 times higher (42% vs 8%) (Figure 3E). These findings suggest that the harsh chemical and extractive treatments involved in melanin ghost preparation are more effective at removing cellular constituents such as polysaccharides than lipids, resulting in the relative enrichment of the latter constituents.

**Figure 3.**
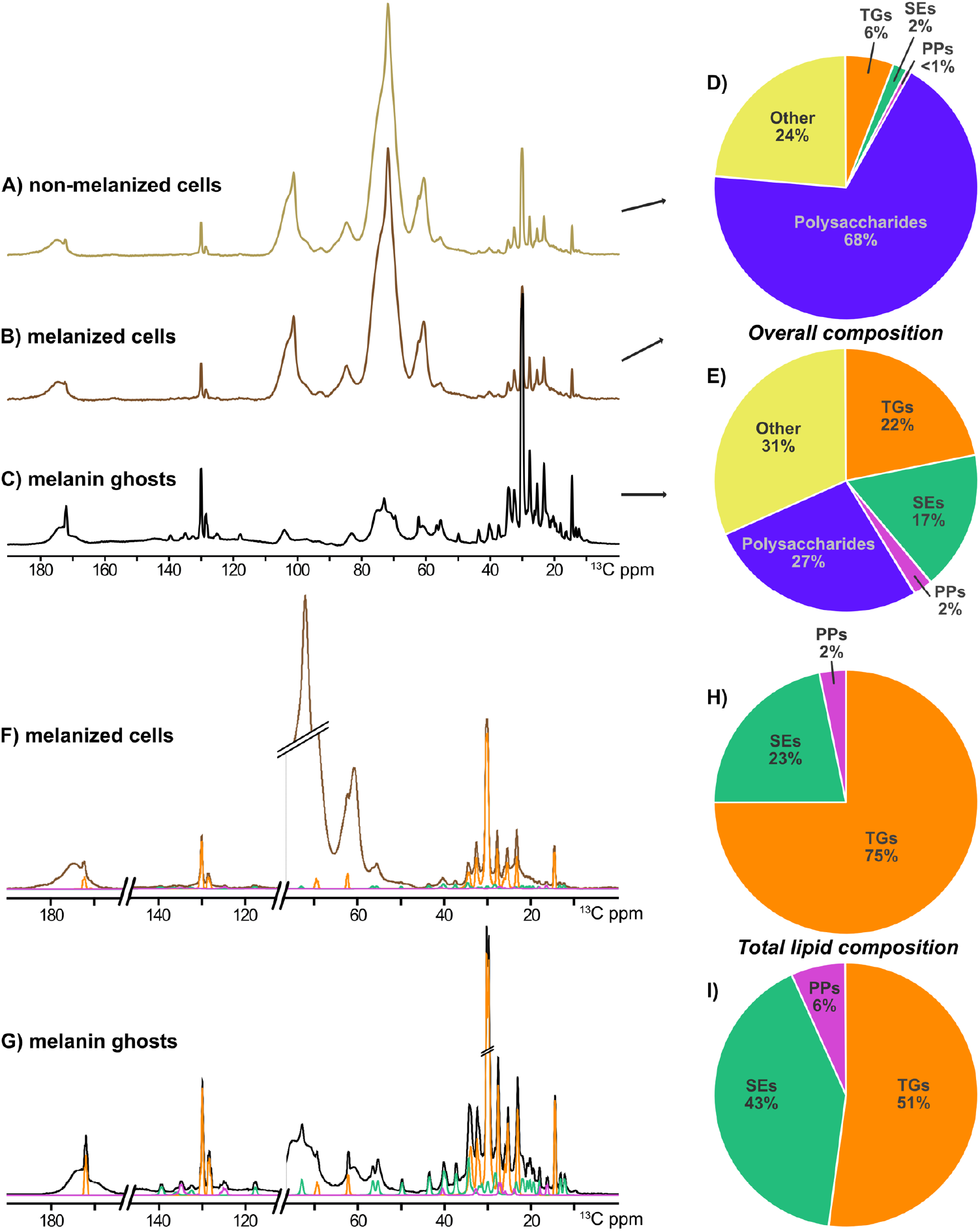
Quantitative estimates of the relative amounts of various cellular components in *C. neoformans* whole cell and melanin ghost samples prepared from cultures containing [U-^13^C_6_]-glucose-enriched media, demonstrating the preferential retention of lipids by melanin ghosts. Top left: ^13^C quantitatively reliable DP (50-s recycle delay) spectra of A) non-melanized and B) melanized whole cells from the acapsular mutant Cap59 strain and C) melanin ghosts generated from H99 cells. Top right: Pie charts displaying the relative quantities of lipid, polysaccharide, and ‘Other’ (unidentified) cellular constituents with respect to their total content in D) whole cells and E) melanin ghosts. Only one pie chart is shown for both melanized and non-melanized whole cells due to the absence of compositional differences between the two samples. Bottom left: ^13^C DP spectra of F) melanized whole cells and G) melanin ghosts overlaid with the fitted curves (obtained with DMfit spectral deconvolution software) that correspond to the three classes of identified lipids. Bottom right: Pie charts displaying the relative quantities of TGs, SEs, and PPs with respect to the total lipid content in whole cells (H) and melanin ghosts (I). Triglycerides, **TG**; orange. Sterol esters, **SE**; green. Polyisoprenoids, **PP**; violet.

Considerable quantitative variations in the relative contributions of TGs, SEs, and PPs that comprise the total lipid content of each sample were also observed between the melanin ghost and whole cell preparations. With respect to the total signal intensity across the spectrum, TGs accounted for the largest signal intensity (22% vs. 6% for melanin ghosts and whole cells, respectively), followed by SEs (17% vs. 2%), and then PPs (2% vs. <1%) (Figures 3D and 3E). With respect to the signal intensity attributable to lipids, the balance among the various lipid species differed substantially between the two types of samples. SEs and PPs corresponded to a much smaller proportion of the total lipid content in whole cells (23% and 2%) as compared with melanin ghosts (43% and 6%, respectively) (Figures 3H and 3I). Interestingly, the constituents that are present in the lowest proportions in whole cells are preferentially retained by melanin ghosts: the SE content of melanin ghosts was increased by 2-fold (23% vs. 43%) and the PP content by 3-fold (2% vs. 6%). Taken together, these data suggest that in spite of the lack of influence melanin synthesis has on the types or relative amounts of lipids produced by *C. neoformans* cells, melanin pigments preferentially associate with certain lipid species, namely TGs, SEs, and PPs.

## Discussion

One of the most striking features in the ^13^C ssNMR spectra of *C. neoformans* melanin ghosts is the prominent group of sharp upfield peaks (~10-40 ppm) that are characteristic of lipid long-chain methylene carbons. The quantitative ssNMR analyses conducted previously and in the current study have estimated that lipids comprise 40-70% of the total NMR signal intensity in ^13^C DPMAS melanin ghost spectra, whereas polysaccharides account for at most 30% (13). It is now well-established that the cell-wall polysaccharides, in particular chitin and its deacetylated derivative chitosan, play a key role in anchoring *C. neoformans* melanin pigments to the cell wall. Genetic disruption of the chitin/chitosan biosynthetic machinery compromises cell-wall integrity and has deleterious effects on pigment retention, resulting in a “leaky melanin” phenotype (14, 15). Conversely, supplementing *C. neoformans* growth cultures with GlcNAc, the monomeric building block of chitin, increases cell-wall melanin pigment deposition and retention (43). Thus, it is unsurprising in retrospect that chitinous polysaccharides are retained in ghosts due to their intimate association with the melanin pigment. However, there is no currently proposed rationale for the presence of substantial quantities of lipids in melanin ghosts.

In this work, we set out to identify the particular lipid molecular classes that are strongly associated with the melanin pigment to begin the process of unraveling the roles of these lipids in *C. neoformans* melanization. The implementation of 2D ssNMR experiments that preferentially detect the signals of mobile molecular constituents allowed us to uncover TGs, SEs, and PPs as the primary lipids in uniformly ^13^C-enriched melanin ghost samples. Although prior work by our group has provided evidence for the presence of TGs (18), determined herein to be the most abundant lipid species, SEs and PPs have not been reported previously as melanin ghost constituents.

The fact that no lipid species other than TGs, SEs, and PPs were observed in our ghost samples also strongly suggests that these constituents originate from lipid droplets (LDs), subcellular organelles that function as storage depots for neutral lipids in all living organisms (44). It is well established that LDs are enclosed by a phospholipid monolayer and that TGs and SEs comprise approximately 95% of their contents (45, 46). Although PPs are not considered to be typical LD constituents, their presence in these organelles is not unprecedented: PPs have been detected in LDs isolated from *Saccharomyces cerevisiae* (47).

All organisms produce PPs, namely polyprenols and dolichols, which function in the endoplasmic reticulum (ER) as sugar carriers for reactions such as protein glycosylation and GPI anchor synthesis (48, 49). PPs have also been found to accumulate in a vast array of other organelles and tissue types, but the roles they play in these cellular locations are currently uncertain (50). Nonetheless, PPs within the LDs of *S. cerevisiae* have been proposed to function by coordinating the assembly of the cell wall of sporulating fungal cells: the delivery of PPs to the cell wall via LDs is thought to activate chitin synthase 3 (Chs3), resulting in the formation of cell-wall chitin that is subsequently deacetylated to chitosan (47). Thus, it is informative to consider our findings for key polysaccharide and lipid species in their cellular context: 1) melanin ghosts are essentially isolates of the melanized cell wall; 2) chitin and chitosan play a critical role in the anchoring of melanin pigments to the *C. neoformans* cell wall; 3) both of these polysaccharides, along with PPs, TGs, and SEs, have been identified within melanin ghosts; 4) TGs, and SEs are known to be typical LD cargo. Together, these considerations suggest that PPs could also be contained within *C. neoformans* LDs, where they would play analogous roles to the PPs reported in *S. cerevisiae*.

The exceptionally sharp TG, SE, and PP resonances in our ^13^C ssNMR spectra of melanin ghosts and whole *C. neoformans* cells indicate that these lipids continue to undergo the rapid isotropic motions typical of small molecules in solution despite being present in the solid state. Unlike the amphiphilic lipids of cell membranes that adopt layered arrangements in aqueous environments, neutral lipids such as TGs and SEs lack the charged functional groups that are required to form ordered structures and can therefore retain a significant amount of molecular mobility even when packed in LDs (44). Given the prior micrographic evidence that *C. neoformans* cells contain abundant intracellular stores of LDs (51–54), we can suggest that at least a portion of the exceptionally narrow lipid signals displayed by the whole cells arise from TGs, SEs, and PPs that are contained within LDs. However, *C. neoformans* LDs have not been found to localize to the cell wall; they have only been observed in the cytoplasm (51, 52), the contents of which are removed during melanin ghost isolation. Thus, it is unlikely that the lipids retained by ghosts remain packed inside these organelles. Instead, we propose that a portion of the lipids are released from LDs and transferred to the cell wall, possibly in conjunction with melanin pigment deposition or during other cell wall transport processes (12). Consequently, a subset of these LD-derived lipids become trapped within the cellular framework of the melanized cell wall. Nonetheless, rather than becoming immobilized by association or covalent bonding to the cell wall or pigment, the lipids are likely to be contained within discrete cellular domains that allow them to undergo the molecular motions that produce sharp ssNMR spectral lines. Although this a preliminary hypothesis that requires rigorous experimental testing, our findings nevertheless argue in favor of a role for LDs in *C. neoformans* melanization.

Surprisingly, the lipid profile we determined for melanin ghosts is not in accord with prior reports of *C. neoformans* lipid composition. Although these reports include a few particular variations, the discrepancies with respect to our findings are more widespread. First, constituents such as phospholipids and free sterols (54, 55) are notably absent in our melanin ghost spectra. Secondly, the species of SEs we identified (ergosta-7-enol and ergosta-7,22-dienol) had typically been reported as only minor sterol constituents (56, 57) or were absent altogether (58, 59). Third, as noted above there have been no reports of PPs as major lipids produced by *C. neoformans*. Because these prior analyses were carried out exclusively on non-melanized cells, we considered that *C. neoformans* cells might have a distinct lipid profile when grown under melanizing conditions (i.e., in the presence of an obligatory exogenous melanization substrate such as L-DOPA and in minimal media, which encourages melanization), which might well result in a distinctive lipid composition for melanin ghosts. Nevertheless, ssNMR examination of whole acapsular *C. neoformans* cells and melanin ghosts under identical conditions invalidates this hypothesis: our spectroscopic analyses (Figures 3A-I) clearly indicated that melanized and non-melanized cells contain the same classes and relative amounts of lipids.

Because the unexpected anomalies in lipid compositional profile for melanin ghosts (Δ^7^ sterol esters, TGs and PPs) were replicated more generally in both melanized and non-melanized *C. neoformans* cells, it was important to ask why our results varied so significantly from those of other investigators. A degree of variation could be attributed to our choice of methodology: lipid analyses have traditionally been conducted on extracted material using mass spectrometry, whereas our identifications and quantitative estimates are made directly on whole cells. However, preliminary comparisons (Hau-Ming Jan, personal communication) do not support this explanation.

A more compelling hypothesis, however, concerns legitimate but significant differences between the prior and current conditions used for culturing the *C. neoformans* cells. To facilitate isolation of melanin ghosts, our cells were grown under conditions that promote robust melanization in the presence of an appropriate obligatory catecholamine substrate (60–62), namely 10-14 days of culture in a chemically defined minimal medium that contains a low concentration of glucose and only a single nitrogen source. In contrast, all previously published analyses of *C. neoformans* lipids were conducted on cells grown in nutrient-rich media and harvested after significantly shorter incubation times of ~2 days (56–59), leaving open the possibility that lipid composition depends on cellular growth conditions.

A closer examination of the relationships between culture conditions and lipid composition supports the supposition that our incubation of *C. neoformans* cells in nutrient-limited media for exceptionally long time periods can account for profiles that are rich in TGs, SEs, and PPs – and conversely deficient in phospholipids, free sterols, and ergosterol. First, Itoh, et al. (54) analyzed the lipid composition of *C. neoformans* at different phases of cell growth. These researchers found that during the early logarithmic phase (~5 hr), phospholipid levels are high and neutral lipid levels are low; over time, however, the phospholipid content decreases whereas the content of neutral lipids increases. For instance, neutral lipids comprised ~80% of total lipid content for cells harvested at the stationary phase of growth (~71 hr), and the vast majority (>90%) were TGs. A similar trend was observed in relative sterol content: the ratio of sterol esters to free sterols increased from 1:6.9 to 1.3:1 as the cells progressed from early logarithmic to stationary phases of growth. Secondly, work on *C. neoformans* sterol composition by Kim, et al. (63) is the sole report of Δ^7^ sterol ergosta-7-enol as the major species of sterol in *C. neoformans*; notably, it is also the only study carried out on cells harvested after an unusually long 5-day growth period. Thirdly, although we located no evidence for PPs outside of the ER in *C. neoformans* cells, two reports identified PPs in various subcellular locations in the *Aspergillus* species *A. fumigatus* and *A. niger* (64, 65). Both studies were conducted on cells grown for over 9 days, the time requirement determined by Barr et al. (1972) to achieve maximum concentration. Thus, in the context of the available literature, our findings support cell age and nutrient deprivation as modulators of lipid production and composition in *C. neoformans*.

Notably, cell aging and nutrient deprivation have each been linked to pathogenesis in *C. neoformans*. First, cells that have undergone many replicative cycles and are thus older generationally have been shown to accumulate during chronic cryptococcal infections and manifest distinct phenotypic traits associated with virulence (66, 67). For example, these cells display a large body size and thicker cell wall, both of which have been correlated with increased drug resistance and decreased susceptibility to killing by macrophages. Secondly, nutrient deprivation induces several virulence-associated determinants, for example, by increasing the gene expression and activity of the laccase enzyme Lac1 (68, 69). Since this laccase catalyzes the rate-limiting step in melanin biosynthesis, one way in which it contributes to virulence is by initiating pigment formation (assuming the availability of melanization substrate) (70, 71). Additionally, Lac1 plays a key role in other biosynthetic pathways that yield virulence-associated products (72), e.g., prostaglandin (PG) signaling lipids (73) produced from polyunsaturated fatty acids (PUFAs) (74). Specifically, Lac1 is essential for the formation of the “authentic” PGs prostaglandin-E2 (PGE2) and 15-keto-prostaglandin-E2 (15-keto-PGE2) (75, 76), which are largely produced by vertebrates to regulate immunomodulatory and inflammatory processes (77, 78). Indeed, the synthesis of these PGs by *C. neoformans* during *in vivo* infection has been found to be required for cell growth and proliferation within macrophages (79). Considering that Lac1 expression is induced by nutrient deprivation even in the absence of melanization substrate, and that the vast majority of lipids in both the melanized and non-melanized cells we examined were identified as PUFAs in the form of TGs, it is possible that nutrient deprivation specifically increases the production of these lipids, which are then stored in LDs as readily available precursors for PG biosynthesis. In sum, it is likely that the lipid species we detected in cells grown for a prolonged period of time in nutrient-limited media are the predominant lipid species produced during *in vivo* infection and could contribute to virulence independently of melanin formation.

Interestingly, the impact of cell age and cellular stress on PP production has been documented in a diverse range of organisms, although not commonly in fungi. A key example is the accumulation of the PP dolichol in various human tissues: the quantity of this lipid increases by several orders of magnitude throughout the human lifespan in tissues such as the liver, thyroid, and testis (80), and in fact it has been proposed as a biomarker of cellular aging (81). The contention that accumulation of cellular PP is a general mitigating response to cellular stress (82) is supported by evidence in both mammals and plants. For example, dolichols act as radical scavengers of peroxidized lipids and could thereby reduce the oxidative damage associated with the progression of age-related mammalian diseases (83). PPs such as polyprenol and solanesol accumulate within plant leaves in response to both light and viral infections (84, 85). Despite the relative paucity of reports linking PP levels to stress in fungi, it is worth noting that the isopentenyl diphosphate isomerase enzyme of the isoprenoid pathway is the product of an essential gene in fungal species such as *S. cerevisiae* (86), *Schizosaccharomyces pombe* (87), and *C. neoformans* (88); conditional mutation of this gene results in cells with reduced stress tolerance (89). Given this body of evidence, it is plausible that in our *C. neoformans* cellular system, the stress induced by a nutrient-limited environment for a prolonged period of time (i.e., MM for 10 days) elevates the production of PPs.

Although we found no variations in lipid species between melanized and non-melanized *C. neoformans* cells, there is an interesting parallel between the lipids we identified and those found associated with mammalian neuromelanin: dolichol and its derivatives are well-established components of neuromelanin (90, 91). Moreover, TEM analysis has demonstrated that these “melanic-lipidic” granules are contained in organelles complexed together with LDs (92, 93). Analogous to the accumulation of dolichol and its hypothesized role as a protective compound, neuromelanin builds up in the brain tissue of all mammals with age and is thought to protect against neuronal degeneration by sequestering reactive neurotoxic substrates (94). In contrast, no rationale has been proposed for the presence of LDs in the neuromelanin literature. However, there is evidence to suggest that in endolichenic fungal species, the trapping of lipophilic toxins by LDs is a so-called self-resistance mechanism that quenches the production of reactive oxygen species and thereby contributes to the ability of these fungi to live among algae, cyanobacteria, or other photosynthesizing microorganisms in a mutualistic relationship (95). Since melanin formation is an oxidative process that involves the generation of multiple reactive intermediates (96–98), it is plausible that in melanotic organisms, the LDs located at sites where melanin synthesis takes place could have a similar function. The fact that lipids originating from LDs were also predominant in the non-melanized *C. neoformans cells* could additionally suggest that they are produced in anticipation of melanin synthesis: the cell culture conditions that promote melanin synthesis could simultaneously promote the synthesis of LDs in preparation for acquiring a substrate suitable for melanization.

In summary, the work herein provides a comprehensive compositional analysis of the lipids that associate with the melanin pigments produced by the fungal pathogen *C. neoformans*, also exploring their possible origins and roles in cellular function. By implementing solid-state NMR strategies that display resonances from mobile molecular species preferentially, we bypassed the need for traditional lipid extraction. We were able to examine solid samples of the melanized cell wall directly under near-native conditions, uncovering triglycerides, sterol esters, and polyisoprenoids as the major lipid components and estimating their relative proportions. In turn, our findings suggested a role for lipid droplets in the melanization process and a dependence of lipid composition on cell culture conditions. In addition, we compared the NMR spectra of intact cells grown under melanizing and non-melanizing conditions, determining the lipid composition of melanized *C. neoformans cells* for the first time. Our results indicate an intimate association between lipids and melanin pigment that must be accounted for in proposed schema for fungal melanization and cell wall melanin architecture.

## Experimental procedures

### *C. neoformans* strains and cell growth

The well-characterized serotype A strain H99 was used for melanin ghost isolation and the acapsular mutant strain Cap59 (generated from an H99 background (99)) was used for whole cell ssNMR analysis. Both isolates were maintained in 20% glycerol stock at −80 °C. The cells were inoculated into Sabouraud dextrose broth and grown for 48 h at 30 ̊C with moderate shaking (120 rpm) prior to harvest.

### Culture conditions

The harvested cells grown in Sabouraud dextrose broth were washed with Phosphate-Buffered Saline (PBS), counted using a hemocytometer, and adjusted to a cell concentration of 1×10^6^ mL^−1^ in minimal media (MM) containing ^13^C-enriched glucose as the sole carbon source (29.4 mM KH_2_PO_4_, 10 mM MgSO_4_, 13 mM glycine, 3 μM thiamine, 15 mM [U-^13^C_6_]-glucose, pH 5.5). The absence of an obligatory exogenous melanization precursor in this media recipe resulted in the production of non-melanized yeast cells. To produce melanized cells, 1 mM L-DOPA was added to the MM formulation listed above. All cultures were grown for 10 days at 30 °C with shaking at moderate speed (120 rpm). The cells were collected by centrifugation, resuspended in PBS, and stored at −20 °C until further use.

### Isolation of melanin ‘ghosts’

*neoformans* H99 cells cultured in MM containing [U-^13^C_6_]-glucose and 1 mM L-DOPA for 10 d were subjected to the standard melanin ghost isolation protocol as described in our prior studies (17). Briefly, the cells obtained from 100 mL of a melanized culture were subjected to enzymatic hydrolysis of cell-wall and protein constituents, followed by three consecutive Folch lipid extractions, and then boiling in 6 M HCl for 1 hr. The resulting black solid particles were obtained by centrifugation, dialyzed against distilled water for several days, and subsequently lyophilized for 3 days.

### Preparation of whole fungal cells

Cultures of melanized and non-melanized *C. neoformans* cells from the acapsular mutant strain Cap59 were grown side by side in separate flasks containing [U-^13^C_6_]-glucose-enriched MM using the same culture conditions described above. L-DOPA was added to achieve a concentration of 1 mM in one of the flasks to produce melanized cells. The cell pellet obtained from a 100 mL aliquot of each culture was resuspended in 25 mL of deionized water, vortexed vigorously, and then centrifuged at 17,000 rpm for 20 min at 4 °C. The supernatant was decanted, and this process was repeated four more times to remove any residual metabolites or other small molecules. After the fifth wash, the cell pellet was lyophilized for 3 days prior to analysis by ssNMR.

### Solid-State NMR measurements

Solid-state NMR experiments reported in this work were carried out on a Varian (Agilent) DirectDrive2 (DD2) instrument operating at a ^1^H frequency of 600 MHz and equipped with a 3.2-mm T3 HXY MAS probe (Agilent Technologies, Santa Clara, CA). All data were acquired on ~12-18 mg of lyophilized melanin ghost or whole cell samples using a Magic Angle Spinning (MAS) rate of 15.00 ± 0.02 kHz and a spectrometer-set temperature of 25 °C. The two-dimensional (2D) double-quantum (DQ) ^13^C-^13^C correlation spectra were measured using the refocused J-INADEQUATE pulse sequence (100, 101), which relies on scalar (J) coupling for polarization transfer between directly bonded carbon nuclei. The experiments were carried out with ^13^C 90° and 180° pulse lengths of 2.6 μs and 5.2 μs, respectively, a ^1^H 90° pulse length of 2.3 μs, and 2.4 ms τ spin echo delays. Direct polarization (DP) was used for initial ^13^C excitation and a 2-s recycle delay was implemented to enhance the signals of mobile constituents. Small phase incremental alternation pulse sequence (SPINAL) decoupling (102) was applied during acquisition at a radio frequency (rf) field of 109 kHz. The 2D ^1^H-^13^C correlation spectra were measured with a solid-state HETCOR pulse sequence that used refocused INEPT for the selective transfer of polarization between directly bonded proton-carbon pairs located within mobile molecules (20). The experiments were carried out with ^13^C and ^1^H 90° pulse lengths of 2.5 μs and 2.3 μs, respectively, and a 2-s recycle delay. Inter-pulse delays were optimized for maximum ^1^H-^13^C transfer efficiency using time periods of 1/4J and 1/6J (with a ^1^H-^13^C coupling constant of J = 145 Hz), corresponding to 1.72 ms and 1.15 ms, respectively. ^1^H decoupling using the WALTZ method (103) with an rf field of 5.5 kHz was applied during signal acquisition. The one-dimensional (1D) ^13^C DPMAS experiments used to estimate the relative proportions of carbon-containing cellular constituents in melanin ghost and whole cell samples were implemented with long recycle delays (50 s) to obtain quantitatively reliable signal intensities. The ^13^C and ^1^H 90° pulse lengths were 2.6 μs and 2.3 μs, respectively. SPINAL decoupling with an rf field of 109 kHz was applied during acquisition.

### Solid-State NMR quantitative analysis

Spectral curve fitting was performed with the deconvolution software DMfit (104) by inputting the ^13^C frequency and linewidth of each peak identified in the J-INADEQUATE spectra and measuring their individual integrated intensities. The relative quantities of lipid, polysaccharide, and ‘Other’ cellular constituents were estimated by expressing the sum of the individual peak areas corresponding to each constituent type as a fraction of the total integrated area of the NMR spectrum. The relative composition of triglycerides, sterol esters, and polyisoprenoids was estimated by expressing the summed peak areas as a fraction of the total integrated signal intensity attributed to lipids. The fitted curves of peaks located in crowded spectral regions were adjusted manually to match their integrated areas with the average from a set of fully resolved peaks corresponding to each constituent.

## Data availability

The ssNMR datasets presented in this study and the pulse sequences implemented to acquire them are available from the corresponding author upon request.

## Acknowledgements

We thank Ankur Jadhav (Chemical Engineering, City College of New York) for his expert instruction and ongoing support in the use of the software DMfit. We also thank Van Chanh Phan (Hostos Community College, CUNY) for his development of ssNMR programs and technical support with experimental setup.

## Funding and additional information

This work was supported by National Institutes of Health Grant R01-AI052733. The 600 MHz NMR facilities used in this work are operated by The City College (CCNY) and the CUNY Institute for Macromolecular Assemblies. C.C. was the recipient of a fellowship award from the U.S. Department of Education Graduate Assistance in Areas of National Need (GAANN) Program in Biochemistry, Biophysics, and Biodesign at The City College of New York (PA200A120211 and PA200A150068) and the James Whittam award of the CCNY Department of Chemistry & Biochemistry.

## References

1. Rajasingham, R., Smith, R. M., Park, B. J., Jarvis, J. N., Govender, N. P., Chiller, T. M., Denning, C. W., Loyse, A., and Boulware, D. R. (2017) Global burden of disease of HIV-associated cryptococcal meningitis: an updated analysis. Lancet Infect. Dis. 17, 873–881

2. Goldman, D. L., Khine, H., Abadi, J., Lindenberg, D. J., Pirofski, L., Niang, R., and Casadevall, A. (2001) Serologic Evidence for Cryptococcus neoformans Infection in Early Childhood. Pediatrics. 107, e66

3. Denham, S. T., and Brown, J. C. S. (2018) Mechanisms of pulmonary escape and dissemination by Cryptococcus neoformans. J. Fungi. 4, 1–17

4. Wozniak, K. L., Olszewski, M. A., and Wormley, F. L. (2015) Molecules at the interface of Cryptococcus and the host that determine disease susceptibility. Fungal Genet. Biol. 78, 87–92

5. Zaragoza, O. (2019) Basic principles of the virulence of Cryptococcus. Virulence. 10, 490–501

6. Eisenman, H. C., and Casadevall, A. (2012) Synthesis and assembly of fungal melanin. Appl. Microbiol. Biotechnol. 93, 931–940

7. Nosanchuk, J. D., Valadon, P., Feldmesser, M., and Casadevall, A. (1999) Melanization of Cryptococcus neoformans in Murine Infection. Mol. Cell. Biol. 19, 745–750

8. Rosas, Á. L., Nosanchuk, J. D., Feldmesser, M., Cox, G. M., McDade, H. C., and Casadevall, A. (2000) Synthesis of polymerized melanin by Cryptococcus neoformans in infected rodents. Infect. Immun. 68, 2845–2853

9. Zaragoza, O., Rocio, G.-R., Nosanchuk, J. D., Cuenca-Estrella, M., Rodriguez-Tudela, J. L., and Casadevall, A. (2010) Fungal Cell Gigantism during Mammalian Infection. PLoS Pathog. 6, 1–18

10. Buchanan, K. L., and Murphy, J. W. (1998) What Makes Cryptococcus neoformans a Pathogen? Emerg. Infect. Dis. 4, 71–83

11. Nosanchuk, J. D., and Casadevall, A. (2006) Impact of melanin on microbial virulence and clinical resistance to antimicrobial compounds. Antimicrob. Agents Chemother. 50, 3519–3528

12. Camacho, E., Vij, R., Chrissian, C., Prados-Rosales, R., Gil, D., O’Meally, R. N., Cordero, R. J. B., Cole, R. N., McCaffery, J. M., Stark, R. E., and Casadevall, A. (2019) The structural unit of melanin in the cell wall of the fungal pathogen Cryptococcus neoformans. J. Biol. Chem. 294, 10471–10489

13. Chrissian, C., Camacho, E., Fu, M. S., Prados-Rosales, R., Chatterjee, S., Cordero, R. J. B. X., Lodge, J. K. X., Casadevall, A., and Stark, R. E. (2020) Melanin deposition in two Cryptococcus species depends on cell-wall composition and flexibility. J. Biol. Chem. 295, 1815–1828

14. Baker, L. G., Specht, C. A., Donlin, M. J., and Lodge, J. K. (2007) Chitosan, the Deacetylated Form of Chitin, Is Necessary for Cell Wall Integrity in Cryptococcus neoformans n ◻ †. Eukaryot. Cell. 6, 855–867

15. Banks, I. R., Specht, C. A., Donlin, M. J., Gerik, K. J., Levitz, S. M., and Lodge, J. K. (2005) A Chitin Synthase and Its Regulator Protein Are Critical for Chitosan Production and Growth of the Fungal Pathogen Cryptococcus neoformans. Eukaryot. Cell. 4, 1902–1912

16. Wang, Y., Aisen, P., and Casadevall, A. (1996) Melanin, melanin “ghosts,” and melanin composition in Cryptococcus neoformans. Infect. Immun. 64, 2420–2424

17. Chatterjee, S., Prados-Rosales, R., Itin, B., Casadevall, A., and Stark, R. E. (2015) Solid-state NMR Reveals the Carbon-based Molecular Architecture of Cryptococcus neoformans Fungal Eumelanins in the Cell Wall. J. Biol. Chem. 290, 13779–90

18. Zhong, J., Frases, S., Wang, H., Casadevall, A., and Stark, R. E. (2008) Following Fungal Melanin Biosynthesis with Solid-State NMR: Biopolymer Molecular Structures and Possible Connections to Cell-Wall Polysaccharides. Biochemistry. 7, 4701–4710

19. Wang, T., and Hong, M. (2015) Solid-state NMR investigations of cellulose structure and interactions with matrix polysaccharides in plant primary cell walls. J. Exp. Bot. 67, 503–514

20. Soubias, O., Réat, V., Saurel, O., and Milon, A. (2002) High resolution 2D 1H-13C correlation of cholesterol in model membrane. J. Magn. Reson. 158, 143–148

21. Alexandri, E., Ahmed, R., Siddiqui, H., Choudhary, M. I., Tsiafoulis, C. G., and Gerothanassis, I. P. (2017) High resolution NMR spectroscopy as a structural and analytical tool for unsaturated lipids in solution. Molecules. 22, 1–71

22. Li, J., Vosegaard, T., and Guo, Z. (2017) Applications of nuclear magnetic resonance in lipid analyses: An emerging powerful tool for lipidomics studies. Prog. Lipid Res. 68, 37–56

23. Lie Ken Jie, M. S. F., and Mustafa, J. (1997) High-resolution nuclear magnetic resonance spectroscopy - Applications to fatty acids and triacylglycerols. Lipids. 32, 1019–1034

24. Alemany, L. B. (2002) Using simple 13C NMR linewidth and relaxation measurements to make detailed chemical shift assignments in triacylglycerols and related compounds. Chem. Phys. Lipids. 120, 33–44

25. McKenzie, J. M., and Koch, K. R. (2004) Rapid analysis of major components and potential authentication of South African olive oils by quantitative 13C nuclear magnetic resonance spectroscopy. S. Afr. J. Sci. 100, 349–354

26. Gunstone, F. D. (1990) The 13C-NMR spectra of oils containing γ-linolenic acid. Chem. Phys. Lipids. 56, 201–207

27. Mannina, L., Luchinat, C., Emanuele, M. C., and Segre, A. (1999) Acyl positional distribution of glycerol tri-esters in vegetable oils: A 13C NMR study. Chem. Phys. Lipids. 103, 47–55

28. Cantrell, C. L., Ali, A., Duke, S. O., and Khan, I. (2011) Identification of Mosquito Biting Deterrent Constituents From the Indian Folk Remedy Plant Jatropha curcas. J. Med. Entomol. 48, 836–845

29. Thoss, V., Murphy, P. J., Marriott, R., and Wilson, T. (2012) Triacylglycerol composition of British bluebell (Hyacinthoides non-scripta) seed oil. RSC Adv. 2, 5314–5322

30. Vlahov, G. (1999) Application of NMR to the study of olive oils. Prog. Nucl. Magn. Reson. Spectrosc. 35, 341–357

31. Wilson, W. K., Sumpter, R. M., Warren, J. J., Rogers, P. S., Ruan, B., and Schroepfer, G. J. (1996) Analysis of unsaturated C27 sterols by nuclear magnetic resonance spectroscopy. J. Lipid Res. 37, 1529–1555

32. Ragasa, C. Y., Torres, O. B., Shen, C. C., Mejia, M. G. R., Ferrer, R. J., and Jacinto, S. D. (2012) Chemical constituents of Aglaia loheri. Pharmacogn. J. 4, 29–31

33. Takaishi, Y., Ohashi, T., and Tomimatsu, T. (1989) Ergosta-7,22-dien-3β-ol glycoside from Tylopilus neofelleus. Phytochemistry. 28, 945–947

34. Kanagasabapathy, G., Malek, S. N. A., Kuppusamy, U. R., and Vikineswary, S. (2011) Chemical composition and antioxidant properties of extracts of fresh fruiting bodies of Pleurotus sajor-caju (Fr.) singer. J. Agric. Food Chem. 59, 2618–2626

35. Tuckey, R. C., Nguyen, M. N., Chen, J., Slominski, A. T., Baldisseri, D. M., Tieu, E. W., Zjawiony, J. K., and Li, W. (2012) Human Cytochrome P450scc (CYP11A1) Catalyzes Epoxide Formation with Ergosterol. Drug Metab. Dispos. 40, 436–444

36. Suttiarporn, P., Chumpolsri, W., Mahatheeranont, S., Luangkamin, S., Teepsawang, S., and Leardkamolkarn, V. (2015) Structures of phytosterols and triterpenoids with potential anti-cancer activity in bran of black non-glutinous rice. Nutrients. 7, 1672–1687

37. Tanaka, Y., Sato, H., Kageyu, A., and Tomita, T. (1987) Determination of arrangement of isoprene units in pig liver dolichol by 13C-n.m.r. spectroscopy. Biochem. J. 243, 481–485

38. Misiak, M., Koźmiński, W., Kwasiborska, M., Wójcik, J., Ciepichal, E., and Swiezewska, E. (2009) Complete 1H and 13C signal assignment of prenoI-10 with 3D NMR spectroscopy. Magn. Reson. Chem. 47, 825–829

39. Tanaka, Y., Sato, H., and Kageyu, A. (1982) Structural characterization of polyprenols by 13C-n.m.r. spectroscopy: Signal assignments of polyprenol homologues. Polym. Reports. 23, 1087–1090

40. Tanaka, Y., Mori, M., Ute, K., and Hatada, K. (1990) Structure and Biosynthesis Mechanism of Rubber from Fungi. Rubber Chem. Technol. 63, 1–7

41. Zaragoza, O., and Casadevall, A. (2004) Experimental modulation of capsule size in Cryptococcus neoformans. Biol. Proced. Online. 6, 10–15

42. Eisenman, H. C., Chow, S. K., Tsé, K. K., McClelland, E. E., and Casadevall, A. (2011) The effect of L-DOPA on Cryptococcus neoformans growth and gene expression. Virulence. 2, 329–336

43. Camacho, E., Chrissian, C., Cordero, R. J. B., Lopes, L. L., Stark, R. E., and Casadevall, A. (2017) N-acetylglucosamine affects cryptococcus neoformans cell-wall composition and melanin architecture. Microbiol. (United Kingdom). 163, 1540–1556

44. Czabany, T., Wagner, A., Zweytick, D., Lohner, K., Leitner, E., Ingolic, E., and Daum, G. (2008) Structural and biochemical properties of lipid particles from the yeast Saccharomyces cerevisiae. J. Biol. Chem. 283, 17065–17074

45. Leber, R., Zinser, E., Paltauf, F., Daum, G., and Zellnig, G. (1994) Characterization of lipid particles of the yeast, Saccharomyces cerevisiae. Yeast. 10, 1421–1428

46. Tauchi-Sato, K., Ozeki, S., Houjou, T., Taguchi, R., and Fujimoto, T. (2002) The surface of lipid droplets is a phospholipid monolayer with a unique fatty acid composition. J. Biol. Chem. 277, 44507–44512

47. Hoffmann, R., Grabińska, K., Guan, Z., Sessa, W. C., and Neiman, A. M. (2017) Long-chain polyprenols promote spore wall formation in Saccharomyces cerevisiae. Genetics. 207, 1371–1386

48. Chojnacki, T., and Dallner, G. (1988) The biological role of dolichol. Biochem. J. 251, 1–9

49. Krag, S. S. (1998) The importance of being dolichol. Biochem. Biophys. Res. Commun. 243, 1–5

50. Surmacz, L., and Swiezewska, E. (2011) Polyisoprenoids - Secondary metabolites or physiologically important superlipids? Biochem. Biophys. Res. Commun. 407, 627–632

51. Nolan, S. J., Fu, M. S., Coppens, I., and Casadevall, A. (2017) Lipids affect the Cryptococcus neoformans-macrophage interaction and promote nonlytic exocytosis. Infect. Immun. 85, 1–18

52. Nguyen, L. N., and Nosanchuk, J. D. (2011) Lipid droplet formation protects against gluco/lipotoxicity in Candida parapsilosis: An essential role of fatty acid desaturase Ole1. Cell Cycle. 10, 3159–3167

53. Bubb, W. A., Wright, L. C., Cagney, M., Santangelo, R. T., Sorrell, T. C., and Kuchel, P. W. (1999) Heteronuclear NMR studies of metabolites produced by Cryptococcus neoformans in culture media: Identification of possible virulence factors. Magn. Reson. Med. 42, 442–453

54. Itoh, T., Waki, H., and Kaneko, H. (1975) Changes of Lipid Composition with Growth Phase of Cryptococcus neoformans. Agric. Biol. Chem. 39, 2365–2371

55. Rawat, D. S., Upreti, H. B., and Das, S. K. (1984) Lipid composition of Cryptoccous neoformans. Microbiologica

56. Singh, A., MacKenzie, A., Girnun, G., and Del Poeta, M. (2017) Analysis of sphingolipids, sterols and phospholipids in human pathogenic Cryptococcus strains. J. Lipid Res. 58, 2017–2036

57. Lamb, D. C., Corran, A., Baldwin, B. C., Kwon-Chung, J., and Kelly, S. L. (1995) Resistant P45051A1 activity in azole antifungal tolerant Cryptococcus neoformans from AIDS patients. FEBS Lett. 368, 326–330

58. Ghannoum, M. A., Spellberg, B. J., Ibrahim, A. S., Ritchie, J. A., Currie, B., Spitzer, E. D., Edwards, J. E., and Casadevall, A. (1994) Sterol Composition of Cryptococcus neoformans in the Presence and Absence of Fluconazole. Antimicrob. Agents Chemother. 38, 2029–2033

59. Currie, B., Sanati, H., Ibrahim, A. S., Edwards, J. E., Casadevall, A., and Ghannoum, M. A. (1995) Sterol compositions and susceptibilities to amphotericin B of environmental cryptococcus neoformans isolates are changed by murine passage. Antimicrob. Agents Chemother. 39, 1934–1937

60. Nurudeen, T. A., and Ahearn, D. G. (1979) Regulation of melanin production by Cryptococcus neoformans. J. Clin. Microbiol. 10, 724–729

61. Wang, Y., Aisen, P., and Casadevall, A. (1996) Melanin, melanin “ghosts,” and melanin composition in Cryptococcus neoformans. Infect. Immun. 64, 2420–2424

62. Eisenman, H. C., Mues, M., Weber, S. E., Frases, S., Chaskes, S., Gerfen, G., and Casadevall, A. (2007) Cryptococcus neoformans laccase catalyses melanin synthesis from both D- and L-DOPA. Microbiology. 153, 3954–3962

63. Kim, S. J., Kwon Chung, K. J., Milne, G. W. A., Hill, W. B., and Patterson, G. (1975) Relationship between polyene resistance and sterol compositions in Cryptococcus neoformans. Antimicrob. Agents Chemother. 7, 99–106

64. Stone, K. J., Butterworth, P. H., and Hemming, F. W. (1967) Characterization of the hexahydropolyprenols of Aspergillus fumigatus Fresenius. Biochem. J. 102, 443–455

65. Barr, R. M., and Hemming, F. W. (1972) Polyprenols of Aspergillus niger. Biochem. J. 126, 1193–1202

66. Bouklas, T., and Fries, B. C. (2015) Aging as an emergent factor that contributes to phenotypic variation in Cryptococcus neoformans. Fungal Genet. Biol. 78, 59–64

67. Bouklas, T., Pechuan, X., Goldman, D. L., Edelman, B., Bergman, A., and Friesa, B. C. (2013) Old Cryptococcus neoformans cells contribute to virulence in chronic cryptococcosis. MBio. 4, 1–10

68. Pukkila-Worley, R., Gerrald, Q. D., Kraus, P. R., Boily, M. J., Davis, M. J., Giles, S. S., Cox, G. M., Heitman, J., and Alspaugh, J. A. (2005) Transcriptional network of multiple capsule and melanin genes governed by the Cryptococcus neoformans cyclic AMP cascade. Eukaryot. Cell. 4, 190–201

69. Zhu, X., and Williamson, P. R. (2004) Role of laccase in the biology and virulence of Cryptococcus neoformans. FEMS Yeast Res. 5, 1–10

70. Cordero, R. J. B., Camacho, E., and Casadevall, A. (2020) Melanization in Cryptococcus neoformans Requires Complex Regulation. MBio. 11, 1–4

71. Lee, D., Jang, E. H., Lee, M., Kim, S. W., Lee, Y., Lee, K. T., and Bahna, Y. S. (2019) Unraveling melanin biosynthesis and signaling networks in cryptococcus neoformans. MBio. 10, 1–21

72. Zhu, X., Gibbons, J., Garcia-Rivera, J., Casadevall, A., and Williamson, P. R. (2001) Laccase of Cryptococcus neoformans is a cell wall-associated virulence factor. Infect. Immun. 69, 5589–5596

73. Erb-Downward, J. R., Noggle, R. M., Williamson, P. R., and Huffnagle, G. B. (2008) The role of laccase in prostaglandin production by Cryptococcus neoformans. Mol. Microbiol. 68, 1428–1437

74. Gabbs, M., Leng, S., Devassy, J. G., and Aukema, H. M. (2015) Advances in Our Understanding of Oxylipins. Am. Soc. Nutr. 6, 513–540

75. Erb-Downward, J. R., and Huffnagle, G. B. (2007) Cryptococcus neoformans produces authentic prostaglandin E2 without a cyclooxygenase. Eukaryot. Cell. 6, 346–350

76. Noverr, M. C., Phare, S. M., Toews, G. B., Coffey, M. J., and Huffnagle, G. B. (2001) Pathogenic Yeasts Cryptococcus neoformans and Candida albicans Produce Immunomodulatory Prostaglandins. Infect. Immun. 69, 2957–2963

77. Lone, A. M., and Taskén, K. (2013) Proinflammatory and immunoregulatory roles of eicosanoids in T cells. Front. Immunol. 4, 1–15

78. Ricciotti, E., and Fitzgerald, G. A. (2011) Prostaglandins and Inflammation. Arterioscler. Thromb. Vasc. Biol. 31, 986–1000

79. Evans, R. J., Pline, K., Loynes, C. A., Needs, S., Aldrovandi, M., Tiefenbach, J., Bielska, E., Rubino, R. E., Nicol, C. J., May, R. C., Krause, H. M., O’Donnell, V. B., Renshaw, S. A., and Johnston, S. A. (2019) 15-keto-prostaglandin e2 activates host peroxisome proliferator-activated receptor gamma (Ppar-γ) to promote cryptococcus neoformans growth during infection. PLoS Pathog. 15, 1–28

80. Carroll, K. K., Guthrie, N., and Ravi, K. (1992) Dolichol: function, metabolism, and accumulation in human tissues. Biochem. Cell Biol. 70, 382–384

81. Parentini, I., Cavallini, G., Donati, A., Gori, Z., and Bergamini, E. (2005) Accumulation of dolichol in older tissues satisfies the proposed criteria to be qualified a biomarker of aging. J. Gerontol. 60, 39–43

82. Bergamini, E., Bizzarri, R., Cavallini, G., Cerbai, B., Chiellini, E., Donati, A., Gori, Z., Manfrini, A., Parentini, I., Signoria, F., and Tamburini, I. (2004) Ageing and oxidative stress: A role for dolichol in the antioxidant machinery of cell membranes? J. Alzheimer’s Dis. 6, 129–135

83. Bergamini, E. (2003) Dolichol: An essential part in the antioxidant machinery of cell membranes? Biogerontology. 4, 337–339

84. Bajda, A., Konopka-Postupolskaa, D., Krzymowska, M., Hennig, J., Skorupinska-Tudek, K., Surmacz, L., Wojcik, J., Matysiak, Z., Chojnacki, T., Skorzynska-Polit, E., Drazkiewicz, M., Patrzylas, P., Tomaszewska, M., Kania, M., Swist, M., Danikiewicz, W., Piotrowska, W., and Swiezewska, E. (2009) Role of polyisoprenoids in tobacco resistance against biotic stresses. Physiol. Plant. 135, 351–364

85. Bajda, A., Chojnacki, T., Hertel, J., Swiezewska, E., Wójcik, J., Kaczkowska, A., Marczewski, A., Bojarczuk, T., Karolewski, P., and Oleksyn, J. (2005) Light conditions alter accumulation of long chain polyprenols in leaves of trees and shrubs throughout the vegetation season. Acta Biochim. Pol. 52, 233–241

86. Mayer, M. P., Hahn, F. M., Stillman, D. J., and Poulter, C. D. (1992) Disruption and Mapping of IDI1, the Gene for Isopentenyl Diphosphate Isomerase in Saccharomyces cerevisiae. Yeast. 8, 743–748

87. Hahn, F. M., and Poulter, C. D. (1995) Isolation of Schizosaccharomyces pombe isopentenyl diphosphate isomerase cDNA clones by complementation and synthesis of the enzyme in Escherichia coli. J. Biol. Chem. 270, 11298–11303

88. Ianiri, G., Boyce, K. J., and Idnurm, A. (2017) Isolation of conditional mutations in genes essential for viability of Cryptococcus neoformans. Curr. Genet. 63, 519–530

89. Watkins, R. A., King, J. S., and Johnston, S. A. (2017) Nutritional Requirements and Their Importance for Virulence of Pathogenic Cryptococcus Species. Microorganisms. 5, 1–20

90. Fedorow, H., Pickford, R., Hook, J. M., Double, K. L., Halliday, G. M., Gerlach, M., Riederer, P., and Garner, B. (2005) Dolichol is the major lipid component of human substantia nigra neuromelanin. J. Neurochem. 92, 990–995

91. Ward, W. C., Guan, Z., Zucca, F. A., Fariello, R. G., Kordestani, R., Zecca, L., Raetz, C. R. H., and Simon, J. D. (2007) Identification and quantification of dolichol and dolichoic acid in neuromelanin from substantia nigra of the human brain. J. Lipid Res. 48, 1457–1462

92. Halliday, G. M., Ophof, A., Broe, M., Jensen, P. H., Kettle, E., Fedorow, H., Cartwright, M. I., Griffiths, F. M., Shepherd, C. E., and Double, K. L. (2005) α-Synuclein redistributes to neuromelanin lipid in the substantia nigra early in Parkinson’s disease. Brain. 128, 2654–2664

93. Engelen, M., Vanna, R., Bellei, C., Zucca, F. A., Wakamatsu, K., Monzani, E., Ito, S., Casella, L., and Zecca, L. (2012) Neuromelanins of Human Brain Have Soluble and Insoluble Components with Dolichols Attached to the Melanic Structure. PLoS One. 7, 1–13

94. Zecca, L., Bellei, C., Costi, P., Albertini, A., Monzani, E., Casella, L., Gallorini, M., Bergamaschi, L., Moscatelli, A., Turro, N. J., Eisner, M., Crippa, P. R., Ito, S., Wakamatsu, K., Bush, W. D., Ward, W. C., Simon, J. D., and Zucca, F. A. (2008) New melanic pigments in the human brain that accumulate in aging and block environmental toxic metals. PNAS. 105, 17567–17572

95. Chang, W., Zhang, M., Zheng, S., Li, Y., Li, X., Li, W., Li, G., Lin, Z., Xie, Z., Zhao, Z., and Lou, H. (2015) Trapping toxins within lipid droplets is a resistance mechanism in fungi. Sci. Rep. 5, 1–11

96. Ito, S. (2006) Encapsulation of a reactive core in neuromelanin. PNAS. 103, 14647–14648

97. Solano, F. (2014) Melanins: Skin Pigments and Much More—Types, Structural Models, Biological Functions, and Formation Routes. New J. Sci. 2014, 1–28

98. Zucca, F. A., Segura-Aguilar, J., Ferrari, E., Muñoz, P., Paris, I., Sulzer, D., Sarnae, T., Casella, L., and Zecca, L. (2017) Interactions of iron, dopamine, and neuromelanin pathways in brain aging and parkinson’s disease. Physiol. Behav. 155, 96–119

99. Chang, Y. C., and Kwon-Chung, K. J. (1994) Complementation of a capsule-deficient mutation of Cryptococcus neoformans restores its virulence. Mol. Cell. Biol. 14, 4912–4919

100. Bax, A., Freeman, R., and Kempsell, S. P. (1980) Natural Abundance 13C-13C Coupling Observed via Double-Quantum Coherence. J. Am. Chem. Soc. 102, 4849–4851

101. Lesage, A., Auger, C., Caldarelli, S., and Emsley, L. (1997) Determination of through-bond carbon-carbon connectivities in solid-state NMR using the INADEAUATE experiment. J. Am. Chem. Soc. 119, 7867–7868

102. Fung, B. M., Khitrin, A. K., and Ermolaev, K. (2000) An Improved Broadband Decoupling Sequence for Liquid Crystals and Solids. J. Magn. Reson. 142, 97–101

103. Shaka, A. J., Keeler, J., and Freeman, R. (1983) Evaluation of a new broadband decoupling sequence: WALTZ-16. J. Magn. Reson. 53, 313–340

104. Massiot, D., Fayon, F., Capron, M., King, I., Le Calvé, S., Alonso, B., Durand, J. O., Bujoli, B., Gan, Z., and Hoatson, G. (2002) Modelling one- and two-dimensional solid-state NMR spectra. Magn. Reson. Chem. 40, 70–76

105. Ngamskulrungroj, P., Chang, Y., Roh, J., and Kwon-Chung, K. J. (2012) Differences in nitrogen metabolism between cryptococcus neoformans and C. gattii, the two etiologic agents of cryptococcosis. PLoS One. 7, 1–11

106. Kikuchi, G. (1973) The glycine cleavage system: composition, reaction mechanism, and physiological significance. Mol. Cell. Biol. 1, 169–187

107. Ducker, G. S., and Rabinowitz, J. D. (2017) One-Carbon Metabolism in Health and Disease. Cell Metab. 25, 27–42

108. Nes, W. D., Mccourt, B. S., Zhou, W., Ma, J., Marshall, J. A., Peek, L., and Brennan, M. (1998) Overexpression, Purification, and Stereochemical Studies of the Recombinant (S)-Adenosyl-L-methionine: Δ24(25)-to Δ24(28)-Sterol Methyl Transferase Enzyme from Saccharomyces cerevisiae. Arch. Biochem. Biophys. 353, 297–311

109. Cherniak, R., Morris, L. C., Anderson, B. C., and Meyer, S. A. (1991) Facilitated isolation, purification, and analysis of glucuronoxylomannan of Cryptococcus neoformans. Infect. Immun. 59, 59–64

110. Cherniak, R., O’Neill, E. B., and Sheng, S. (1998) Assimilation of xylose, mannose, and mannitol for synthesis of glucuronoxylomannan of Cryptococcus neoformans determined by 13C nuclear magnetic resonance spectroscopy. Infect. Immun. 66, 2996–2998

